# Evidence for a Nod-like signalling system in cyanobacterial symbiosis with *O. sativa*

**DOI:** 10.64898/2026.07.13.738138

**Authors:** Carmen Sánchez del Solar, Lucía Jiménez-Ríos, Ana Jurado-Flores, José Enrique Frías, Vicente Mariscal, Consolación Álvarez

## Abstract

Symbiotic interactions between plants and nitrogen-fixing microorganisms are essential for sustainable agriculture, yet the molecular mechanisms underlying plant–cyanobacterium symbiosis remain poorly understood. In particular, the nature of the signalling mechanisms mediating partner recognition in associations involving *Nostoc* species is largely unknown. Recent proteomic analyses have identified proteins homologous to *rhizobial* Nod factors biosynthetic enzymes in *Nostoc punctiforme*, suggesting the existence of a Nod-like signalling system. However, the functional role of these components has not been experimentally validated.

Here, we investigate the contribution of *nod*-like biosynthetic and regulatory genes to symbiosis by analysing mutants of *N. punctiforme* affected in genes with homology to *nodB* and *nodD*. Phenotypic characterization revealed that disruption of *nodB*-like genes does not impair free-living growth but affects early stages of plant association and colonization. Specifically, the *nodB1* mutant is impaired in plant association and shows a mild defect in colonization, whereas the *nodB3* mutant exhibits a severe defect in colonization. In contrast, *nodD*-like mutants exhibited altered symbiotic phenotypes, with specific regulators differentially affecting interaction and colonization efficiency in rice (*Oryza sativa*). In particular, mutation of *nodD2* and *nodD3* reduced plant association and severely compromised colonization in *Oryza sativa*, with a more pronounced phenotype in *nodD3* mutant.

Altogether, our results provide genetic evidence supporting the involvement of Nod-like components in cyanobacterial symbiosis and suggest the existence of a regulatory and biosynthetic module contributing to plant colonization. These findings shed new light on the evolution and diversity of symbiotic signalling mechanisms across plant–microbe interactions.

## 1. INTRODUCTION

The establishment of symbiotic interactions between plants and nitrogen-fixing microorganisms represents a key biological process for the development of sustainable agricultural practices, particularly in the context of reducing the reliance on synthetic fertilizers (Álvarez et al., 2023). Among these symbioses, filamentous cyanobacteria of the genus *Nostoc* are remarkable for their ability to associate with a wide diversity of plant lineages, ranging from bryophytes to angiosperms, including economically important crops such as rice (*Oryza sativa*) (Adams et al., 2013; Álvarez et al., 2023). In these interactions, the cyanobacterium provides fixed nitrogen to the host plant, while receiving carbon sources and a protective niche, resulting in enhanced plant growth and productivity (Meeks & Elhai, 2002; Álvarez et al., 2023).

*Nostoc punctiforme* has emerged as a model organism for the study of plant– cyanobacterium symbiosis due to its genetic tractability, ecological versatility, and capacity to establish both extracellular and intracellular associations with multiple plant hosts (Meeks et al., 2001; Álvarez et al., 2023). This filamentous cyanobacterium is capable of differentiating into distinct cell types, including vegetative cells, heterocysts specialized in nitrogen fixation, and hormogonia, which are short, motile filaments that function as infection units during symbiosis (Meeks & Elhai, 2002). The ability to transition between these cellular states underlies its symbiotic competence and adaptability across different environmental and host contexts.

The symbiotic interaction between *N. punctiforme* and plants proceeds through a series of coordinated stages. Initially, a phase of chemical signalling occurs, during which plant-derived compounds such as flavonoids and other exudates induce the differentiation of hormogonia (Cohen & Yamasaki, 2000). These motile filaments exhibit chemotaxis towards the plant, facilitating physical contact and attachment to root tissues (Nilsson et al., 2006). Subsequently, the cyanobacterium establishes a close association with the plant surface, followed by intracellular colonization in compatible hosts such as rice (Álvarez et al., 2020, 2022). These stages are accompanied by extensive transcriptional and proteomic reprogramming in both partners, involving processes related to chemotaxis, cell adhesion, cell wall remodelling, and defence response (Álvarez et al., 2022).

Despite advances in understanding the physiological and molecular changes occurring during *Nostoc*–plant symbioses, the nature of the signalling molecules mediating partner recognition remains largely unknown. In contrast, in well-characterized symbiotic systems such as rhizobium–legume and arbuscular mycorrhizal associations, lipochitooligosaccharides (LCOs), known as Nod and Myc factors, function as key signalling molecules that activate symbiotic responses in the host plant through the common symbiosis signalling pathway (CSSP) (Oldroyd, 2013; Radhakrishnan et al., 2020). These signalling cascades are triggered by the perception of microbial LCOs via plant receptor-like kinases, leading to downstream calcium spiking and transcriptional reprogramming required for microbial accommodation (Oldroyd, 2013; Kim et al., 2019).

Interestingly, recent quantitative proteomic analyses of the early interaction between *N. punctiforme* and *O. sativa* have identified cyanobacterial proteins with homology to rhizobial Nod factors biosynthetic enzymes, including NodB-like polysaccharide deacetylases, NodD-like transcriptional regulators and NodE-like β-ketoacyl synthases (Álvarez et al., 2022). In addition, proteins with similarity to NodO-like calcium-binding factors have also been reported, suggesting the potential existence of a Nod-like signalling system in cyanobacteria (Álvarez et al., 2023). Although the biochemical nature of putative cyanobacterial signalling molecules remains unresolved, these findings raise the possibility that *Nostoc* species may produce LCO-like compounds or functionally analogous signals capable of inducing plant symbiotic pathways.

Consistent with this hypothesis, genetic studies in rice have demonstrated that key components of the CSSP, including the ion channel POLLUX, the calcium/calmodulin-dependent kinase CCaMK, and the transcriptional regulator CYCLOPS, are required for efficient colonization by *N. punctiforme* (Álvarez et al., 2022). These observations support the notion that cyanobacterial symbiosis may rely, at least in part, on conserved signalling modules shared with other intracellular plant–microbe interactions. However, despite the identification of Nod-like proteins and the requirement of the CSSP in the host plant, the functional role of Nod-like genes in *N. punctiforme* has not yet been experimentally validated, and their contribution to symbiotic competence remains unclear.

In rhizobial symbioses, Nod factor biosynthesis and regulation are central to the establishment of a successful interaction with the host plant. Among the key components of this system, NodB and NodD play essential and complementary roles. NodB encodes a polysaccharide deacetylase that catalyses a critical early step in the biosynthesis of lipochitooligosaccharides (LCOs), converting the nonreducing *N*-acetylglucosamine residue of the chitooligosaccharides chain into glucosamine, through the hydrolytic removal of acetyl groups from the chitin oligosaccharide backbone (John et al., 1993). This reaction generates free amino groups that are essential substrates for subsequent enzymatic modifications, such as acylation and the addition of specific substituents that ultimately determine host specificity (Mergaert & Van Montagu, 1997; Oldroyd, 2013). In parallel, NodD proteins function as LysR-type transcriptional regulators that perceive plant-derived flavonoids and activate the expression of nodulation genes, including those involved in LCO biosynthesis (Oldroyd, 2013; Mus et al., 2016). However, multiple NodD homologues are often present in rhizobia, displaying differential regulatory properties, ligand specificities, and target gene sets, and in some cases responding to additional plant- or environment-derived signals (Perret et al, 2000; Masson-Boivin et al., 2009; Oldroyd, 2013). This regulatory diversity allows a more flexible and fine-tuned control of symbiotic gene expression. Overall, this regulatory module establishes a mechanistic link between host signal perception and the production of symbiotic signals in rhizobia.

The identification of Nod-like enzymes in *N. punctiforme* suggests that analogous biosynthetic and regulatory pathways may exist in cyanobacteria (Álvarez et al., 2022). However, the specific roles of these components and their contribution to symbiotic competence remain unclear. In particular, it is currently unknown whether NodB- and NodD-like proteins participate directly in the regulation and production of signalling molecules involved in plant colonization. Thus, despite their potential relevance, no direct genetic evidence has yet demonstrated their functional role in cyanobacterial–plant symbiosis.

Here, we address this gap by investigating the function of Nod-like components in *N. punctiforme* through the analysis of mutants affected in both biosynthetic (*nodB*) and regulatory (*nodD*) genes. By combining phenotypic characterization with symbiotic assays in rice, we provide evidence that NodD-dependent regulatory pathways and NodB-mediated processes contribute to the establishment of the interaction. Our findings support the existence of a Nod-like regulatory and biosynthetic module in cyanobacteria and shed new light on the evolutionary conservation and diversification of symbiotic signalling mechanisms in plant– microbe interactions.

## 2. MATERIALS AND METHODS

### 2.1. Biological material and growth conditions

*N. punctiforme* ATCC 29133 was grown under standard laboratory conditions in BG11(NaNO_3_ as nitrogen source) (Rippka et al. 1979), BG11_0_ (BG11 medium lacking nitrate) and BG11_0_NH_4_ (BG11 medium lacking nitrate and supplemented with 4 mM NH_4_Cl and 8 mM N-tris (hydroxymethyl)methyl-2-aminoethane sulfonic acid (TES)-NaOH buffer, pH 7.5). Cultures were maintained at 28°C under continuous light at an intensity of approximately 50 μE m ² s ¹. Liquid cultures were grown with gentle shaking, while solid media were prepared by supplementing BG11 with 1.5% agar.

*O. sativa L. ssp. indica var. nipponbare* seeds were washed with 80% (v/v) ethanol, rinsed with distilled water and then surface sterilized with 5% (w/v) calcium hypochlorite for 20 min. The seeds were thoroughly washed with sterilized tap water and germinated on wet filter paper in a container. Seedlings were grown hydroponically in controlled environmental chambers at 28°C, with 75% relative humidity and at 12 h light/dark photoperiod.

### 2.2. Identification of Nod-like genes and in silico analysis

Candidate *nodB* and *nodD* homologues in *N. punctiforme* were identified by BLAST searches using Nod protein sequences from *Rhizobium* species as queries. Homologues were selected based on sequence identity, coverage, and the presence of conserved functional domains.

Structural analysis of candidate proteins was performed using available bioinformatic tools, including protein structure prediction, structural alignment and domain prediction platforms. Predicted three-dimensional structures obtained from AlphaFold were used for structural comparisons of candidate proteins with *R. leguminosarum* bv. phaseoli Nod proteins using the super alignment function in PyMOL. Structural similarity was further evaluated by calculating Z-scores using the DALI Protein Structure Comparison Server and TM-scores with the Tm-score Server. Protein domains were identified using the InterPro database.

### 2.3. Generation of mutant strains

The Δ*nodB1,* Δ*nodB2*, Δ*nodB3,* Δ*nodD2* and Δ*nodD3* deletion-insertion mutants were generated by replacing most of the *orf* of the selected genes with a kanamycin resistance cassette (Ω-*npt*) in a PCR-generated DNA fragment containing the 1-Kb regions flanking the mutated *orf* cloned into the suicide plasmid pRL271 (Black et al., 1993). The Ω-*npt* cassette was constructed by amplifying the *npt* gene from pRL278 (Black et al., 1993) and ligating it to the C.S3 cassette (Prentki & Krisch, 1984) previously digested with *Bsa*AI and *Bsa*BI. This procedure eliminated most of the *aad* gene (encoding for resistance to streptomycin and spectinomycin) while preserving the transcriptional terminators and translational stop codons in both ends. The oligonucleotides used for generation of the mutants and Ω-*npt* cassette are listed in Table S1.

In the case of *nodD1*, a single-crossover insertion mutant was generated. For this construct, a truncated version of *nodD1* was generated by amplifying a DNA fragment containing from +27 to +893 of the *orf* and introducing two in-frame stop codons in 5’ end. This construct was cloned into the suicide plasmid pRL278.

The plasmids were transferred to the cyanobacterial parental strain by triparental mating, with plasmids pRL623 and pRK2013 as helper and conjugative plasmids, respectively (Figurski & Helinski, 1979; Elhai et al., 1988, 1997). Exconjugants were identified by the Nm^r^ phenotype.

For selection of the deletion strains, a *sacB*-mediated method for positive selection of double recombinants was used (Cai & Wolk, 1990).

In all cases the genomic structure of the resultant *N. punctiforme* mutant strain was checked by PCR analysis using the oligonucleotides listed in table S1.

### 2.4. Phenotypic characterization of cyanobacterial strains

Phenotypic analysis of wild-type and mutant strains included assessment of filament morphology, cellular structure, and growth. Filament and cell morphology of liquid BG11_0_ cultures were examined using light microscopy. Special attention was given to cellular differentiation processes, including heterocyst formation and hormogonia differentiation, which were evaluated based on morphological criteria. Heterocyst frequency was determined by analysing approximately 100 filaments per strain. For each filament, heterocyst frequency was calculated as the number of heterocysts divided by the total number of cells (vegetative cells and heterocysts) within that filament × 100. The values obtained for individual filaments were used to calculate the mean ± standard error of the mean (SEM) for each strain. Cell morphology was assessed by measuring the transverse (cell width) and longitudinal (cell length) axes of approximately 130 vegetative cells per strain. Cell dimensions are presented as the mean ± SEM. In addition, the aspect ratio, calculated as the ratio of cell length to cell width, was used as an indicator of cell shape, with values close to 1 corresponding to nearly spherical cells and higher values indicating more elongated cells.

Growth was monitored in both liquid and solid media under standard culture conditions. The ability to grow under different nitrogen sources was evaluated after 9 days by spotting 5, 10, 25 and 50 ng of chlorophyll *a* from BG11_0_ liquid cultures of each strain onto solid BG11_0_, BG11 and BG11_0_NH_4_ media. Growth rate was calculated from the exponential phase of growth curves performed in BG110 medium using a Multi-Cultivator MC-1000-OD, under continuous light (50µmol·m^−2^·s^−1^), at 25 °C and under air-sparged conditions. Optical density at 680 was measured every 30 min.

### 2.5. Plant–cyanobacterium interaction assays

For symbiotic assays, surface-sterilized *O. sativa* seeds were germinated for 7 days and transferred to hydroponic system containing BG11_0_ medium. After 4-5 days of acclimation, *N. punctiforme* cultures grown in BG11_0_ medium were added to hydroponic systems at a final concentration of 1 μg chlorophyll *a* mL ¹. Chlorophyll *a* content was measured according to Arnon (1949). Co-culture experiments were carried out under controlled environmental conditions as described above.

### 2.6. Root association kinetics (biofilm formation)

The kinetics of cyanobacterial association to plant roots was assessed by quantifying biofilm formation at different time points (ranging from 1 to 40 days post-inoculation (dpi)). At each time point, roots were collected and processed to measure the amount of associated cyanobacteria.

Association was quantified by measuring chlorophyll *a* content associated with plant roots, as described above, and normalizing values to root biomass. Results were expressed as chlorophyll *a* per milligram of root tissue (Chl · mg ¹ root) ± SEM.

### 2.7. Colonization assays

Plant colonization was assessed at later time points (typically 35 days post-inoculation). All incubations were performed under continuous orbital agitation unless otherwise stated. Roots were washed under running tap water and surface sterilized by two consecutive washes with a surface-sterilizing solution (1% bleach, 0.1% SDS, 0.2% Tween 20) for 4 min to remove external cyanobacteria. Surface-sterilized roots were rinsed with sterile distilled water for 10 min. After washing, roots were washed with BG11 for 5 min. This medium was supplemented with 17.6 mM NaNO_3_ and 2.5 µg/mL amphotericin B and incubated under standard growth conditions and used as a surface-sterilized control.

Root-sterilized fragments were incubated in BG11 supplemented with 2.5 µg/mL amphotericin B under standard growth conditions, and the amount of endophytic *N. punctiforme* was quantified. Colonization levels were determined by measuring chlorophyll *a* content in processed root samples and normalizing to root weight, as described above. Data is shown as mean ± SEM.

### 2.8. Microscopy analysis

Visualization of cyanobacterial colonization and morphological features was performed using light microscopy and, when required, confocal microscopy. Root samples were analysed using a Leica TCS SP2 confocal microscope to assess the extent of colonization and the distribution of cyanobacterial cells within plant tissues. Imaging was carried out using HC PL APO CS ×10 or HCX PL APO ×63 oil immersion (1.4 NA) objectives. Cyanobacterial autofluorescence was detected upon excitation with an argon laser at 488 nm, with emission collected between 650–700 nm. Root lignin autofluorescence was excited using 476 and 488 nm laser lines and detected in the 510–533 nm emission range (Álvarez et al., 2022). Z-stack image series (25–130 optical sections) were reconstructed and processed using ImageJ software version 1.51 j8 (Schneider et al., 2012).

### 2.9. Statistical analysis

Data are presented as mean ± SEM. Statistical significance between wild-type and mutant strains was evaluated using Student’s t-test. Differences were considered significant at P < 0.05. All experiments were performed with multiple biological replicates (typically n ≥ 3), and representative datasets are shown.

## 3. RESULTS

### 3.1. Identification and genomic characterization of *nod*-like genes in *N. punctiforme*

To identify candidate genes involved in a putative Nod-like signalling system, an *in silico* analysis of the *N. punctiforme* genome was performed using homology-based approaches combined with structural characterization. Protein sequences of well-characterized Nod factors biosynthetic and regulatory components from rhizobia were used as queries in BLAST searches against the *N. punctiforme* genome.

Initial screening using rhizobial NodB and NodD sequences identified several candidate homologues displaying moderate sequence similarity and conserved functional domains. Among these, three genes encoding putative NodB-like polysaccharide deacetylases were selected: Npun_F3876, Npun_R4095, and Npun_F4239. These candidates exhibited sequence identities of 31.3%, 32.4%, and 31.6%, respectively, to NodB from *Rhizobium leguminosarum* bv*. phaseoli* (Table 1), consistent with the expected conservation threshold for members of the carbohydrate esterase family 4 (CE4) (Gunawardana, 2019). In addition, the NodB-like protein encoded by Npun_F3876 has been reported to be strongly induced during early interaction with *O. sativa* (Álvarez et al., 2022).

**Table 1.** Sequence similarity analysis of the predicted NodB- and NodD-like proteins from *N. punctiforme* PCC 73102 compared with the NodB and NodD1 proteins of *R. leguminosarum*. Protein length (amino acids), amino acid identity (%), bit score and E-value obtained from UniProt BLASTp are shown. Higher bit scores together with lower E-values indicate stronger sequence similarity. Structural similarity was further assessed using Z-score values obtained from the DALI Protein Structure Comparison Server and TM-score values obtained from Tm-score.

| Gen ID | Protein | Length (aa) | Identity (%) | Score (Bits) | E- value | Z-score | TM-score |
| --- | --- | --- | --- | --- | --- | --- | --- |
| Npun_F3876 | NodB1 | 303 | 31.3 | 217 | 3.5 e-21 | 27.6 | 0.1318 |
| Npun_R4095 | NodB2 | 277 | 32.4 | 231 | 2.4 e-23 | 27.1 | 0.1529 |
| Npun_F4239 | NodB3 | 225 | 31.6 | 220 | 3.6e-22 | 26.7 | 0.2395 |
| Npun_R5691 | NodD1 | 338 | 32.1 | 76 | 0.02 | 20.1 | 0.4147 |
| Npun_F4297 | NodD2 | 313 | 27.3 | 69 | 0.14 | 19.2 | 0.3183 |
| Npun_F6188 | NodD3 | 337 | 37.3 | 63 | 0.77 | 20.6 | 0.5212 |

In parallel, a set of genes encoding putative NodD-like regulators were identified. Five candidates showing homology to LysR-type transcriptional regulators were initially detected, of which three were selected based on sequence conservation and structural similarity: Npun_R5691, Npun_F4297 and Npun_F6188. These proteins displayed sequence identities of 32.1%, 27.3% and 37.3%, respectively, to NodD regulators from *R. leguminosarum* bv*. phaseoli* (Table 1). Notably, Npun_F4297 was found to be upregulated during early stages of plant interaction (Álvarez et al., 2022).

Structural predictions further supported the classification of these candidates, revealing conserved domains characteristic of LysR-type transcriptional regulators in the NodD homologues, as well as catalytic motifs typical of polysaccharide deacetylases in NodB-like proteins (Fig. 1). Structural comparisons using the DALI and TM-score servers yielded contrasting results for the NodB homologues. Although DALI analysis returned high Z-scores (approximately 27), the corresponding TM-scores were very low (TM-score < 0.17 for NodB1 and NodB2), indicating a lack of significant global structural similarity (Zhang & Skolnick, 2004). This apparent discrepancy may reflect the presence of a highly conserved structural domain or local motifs that strongly contribute to the DALI alignment (Holm, 2020), while substantial differences in the overall protein architecture reduce the TM-score. In contrast, the NodD homologues exhibited both high Z-scores and high TM-scores, supporting a conserved overall structure and strong structural similarity among these proteins (Zhang & Skolnick, 2004; Holm, 2020). Furthermore, NodD3 showed a TM-score > 0.50, suggesting a high probability that it shares the same overall fold as previously characterized NodD proteins (Xu & Zhang, 2010).

**Figure 1.**
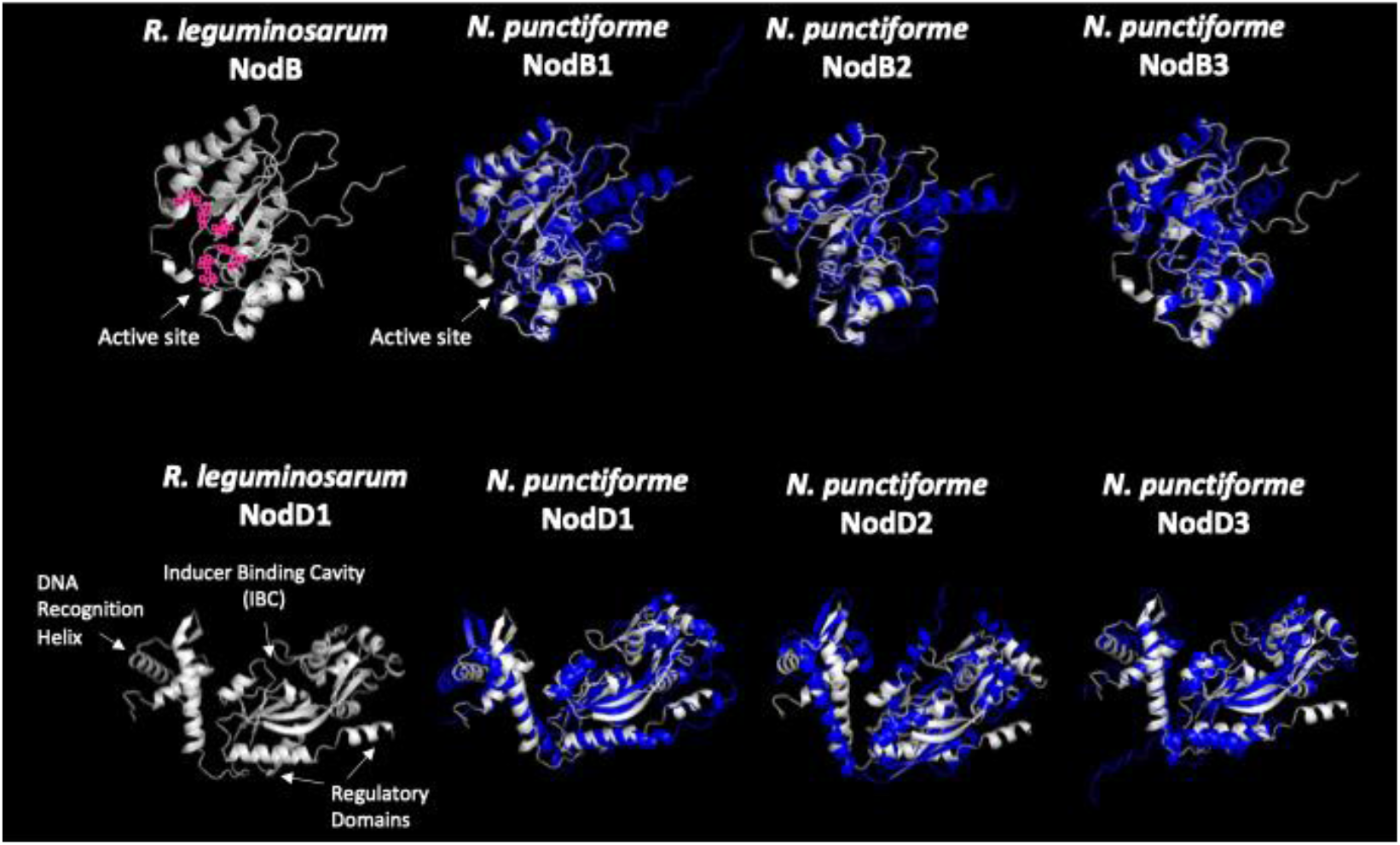
Structural comparison of NodB- and NodD-like proteins from *Nostoc punctiforme*. Structural superposition of the *R. leguminosarum* reference proteins (white) and the corresponding *N. punctiforme* candidate proteins (blue). For NodB, the predicted catalytic residues of the active site are highlighted in pink. For NodD, the DNA-recognition helix, inducer-binding cavity (IBC) and regulatory domains are indicated in the reference structure. Protein superposition was assessed using PyMol. Structures were predicted using AlphaFold and structural superposition was performed using PyMOL (Schrödinger LLC).

Candidate selection was additionally supported by previous in silico studies, including the work of Gunawardana (2019), which identified Npun_R4095 as a putative NodB enzyme based on sequence homology and structural features.

### 3.2. Generation and validation of *nodB* and *nodD* mutant strains

To assess the functional role of Nod-like components in *N. punctiforme*, deletion mutants were generated for selected *nodB*- and *nodD*-like genes. Targeted gene disruptions were performed by double homologous recombination for most candidates, replacing the corresponding open reading frames with an antibiotic resistance cassette. The correct integration of the constructs and the segregation (absence of WT copies) were confirmed at the genomic level by PCR (Fig. S1).

An exception was observed for the *nodD1* mutant. Despite repeated attempts, deletion of this gene by double homologous recombination was not achieved. Therefore, the *nodD1* mutant was generated by single homologous recombination, resulting in insertional disruption of the gene. However, this strain failed to fully segregate, maintaining WT copies in the genome (Supplementary Figure 1).

### 3.3. Disruption of *nodB* does not affect free-living growth but alters plant interaction phenotypes

To determine whether NodB-like enzymes contribute to cyanobacterial physiology under free-living conditions, a detailed phenotypic characterization of the three Δ*nodB* mutants was performed. Filament morphology, cellular structure, and growth were analysed under standard culture conditions. All Δ*nodB* mutants displayed phenotypes indistinguishable from the wild type, with no detectable differences in filament organization, cell morphology, or growth rates (Fig. 2; Table 2), indicating that the selected NodB enzymes are not required for basic cellular functions under these conditions.

**Figure 2.**
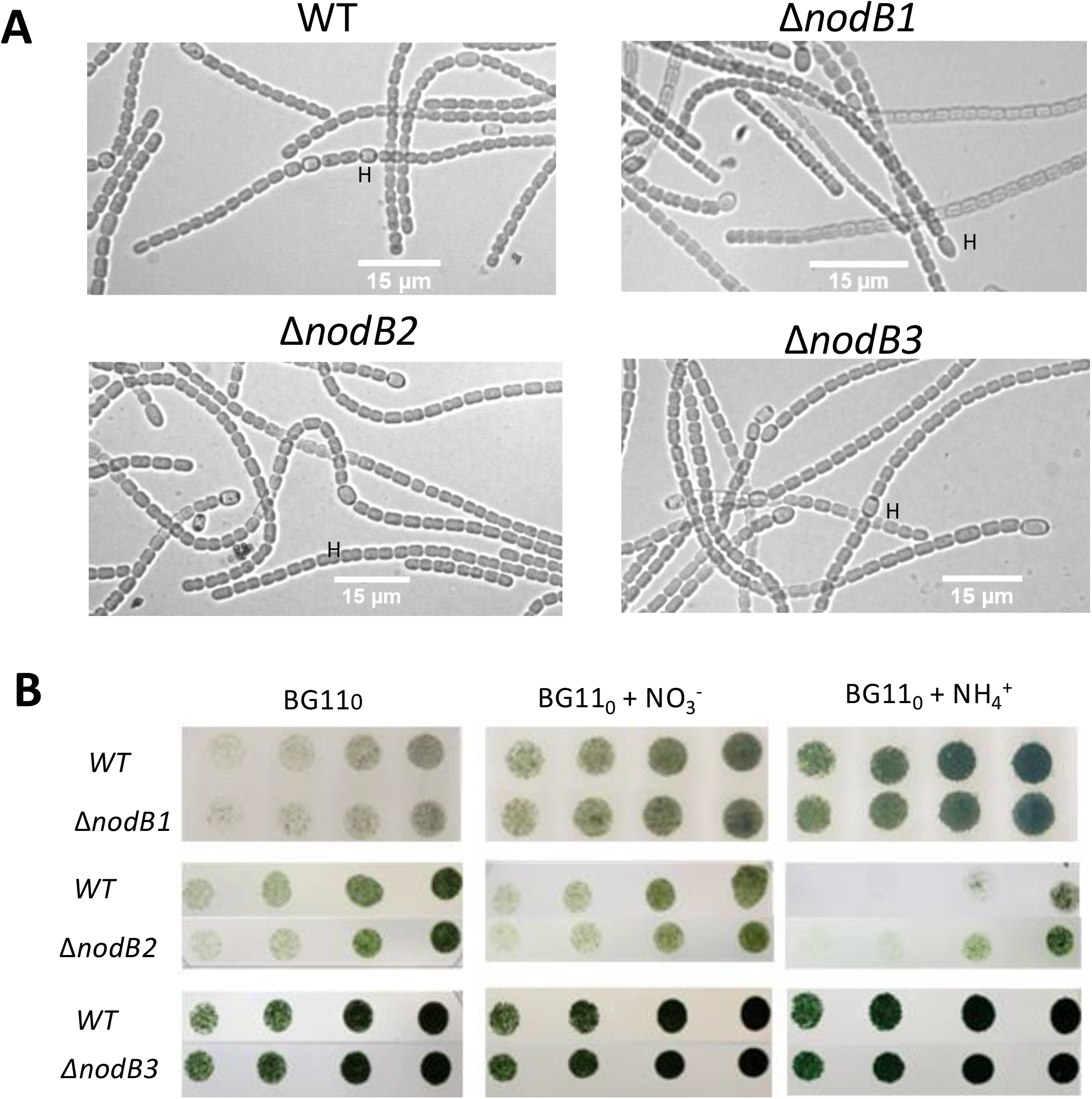
Morphological and growth characterization of *Nostoc punctiforme* WT and *nodB* deletion mutants. (A) Bright-field microscopy images (40× objective) of WT and Δ*nodB1,* Δ*nodB2*, and Δ*nodB3* strains grown in BG110 medium. Vegetative cells and heterocysts are visible; representative heterocysts are indicated with “H”. (B) Spot assay of WT and mutant strains after 9 days of growth on solid media. Serial drops containing 5, 10, 25, and 50 ng of chlorophyll *a* (left to right) were spotted on BG110 and BG110 plates supplemented with combined nitrogen sources, as indicated.

**Table 2.** Morphological characterization of the wild-type (WT) strain and the. Δ*nodB* mutants of *N. punctiforme* grown under diazotrophic conditions. Filament length (number of vegetative cells per filament), heterocyst frequency (% of total cells), vegetative cell length (µm), vegetative cell width (µm), and cell aspect ratio (length/width) are presented as mean ± standard error of the mean (SEM). Hormogonia formation was assessed qualitatively after induction and is indicated as present (+) or absent (−). Quantifications were performed from approximately 100 filaments and 130 cells per condition. Statistical significance is indicated as: ^a^ p < 0.05; ^b^ p < 0.001.

| Strain | Filament length (cells filament<sup>-1</sup>) | Heterocyst percentage (%) | Cell Length (μm) | Cell width (μm) | Aspect ratio (L/W) | Hormogonia formation |
| --- | --- | --- | --- | --- | --- | --- |
| WT | 22.21 ± 0.89 | 6.90 ± 0.38 | 2.50 ± 0.06 | 2.17 ± 0.02 | 1.17 ± 0.03 | + |
| ΔnodB1 | 26.98 ± 1.14<sup>a</sup> | 8.11 ± 0.41<sup>a</sup> | 2.12 ± 0.05<sup>b</sup> | 1.89 ± 0.03<sup>b</sup> | 1.15 ± 0.03 | + |
| ΔnodB2 | 25.17 ± 1.18 | 7.28 ± 0.55 | 2.68 ± 0.05<sup>a</sup> | 2.06 ± 0.02<sup>b</sup> | 1.32 ± 0.03<sup>b</sup> | + |
| ΔnodB3 | 26.70 ± 1.10<sup>a</sup> | 6.70 ± 0.41 | 2.58 ± 0.06 | 2.01 ± 0.03<sup>b</sup> | 1.29 ± 0.03<sup>b</sup> | + |

To evaluate the role of NodB-like proteins during plant interaction, the kinetics of association to *O. sativa* roots was analysed by quantifying biofilm formation over time. Clear differences were observed during the early stages of interaction in the Δ*nodB1* mutant, which exhibited reduced and delayed attachment over time, whereas the Δ*nodB2* and Δ*nodB*3 mutants displayed an overall association profile comparable to that of the wild type, despite isolated differences at specific time points (Fig. 3). At 13–30 days post-inoculation, Δ*nodB1* exhibited a delay in root association, with reductions of approximately 30% compared to the wild type (e.g., 15 dpi: 0.51 ± 0.03 vs 0.34 ± 0.019; P < 0.01; 30 dpi: 0.51 ± 0.023 vs 0.40 ± 0.003; P < 0.01). Although the magnitude of this defect progressively decreased over time, Δ*nodB1* remained impaired in root association, exhibiting an approximately 8% reduction relative to the wild type at 40 dpi (P < 0.05).

**Figure 3.**
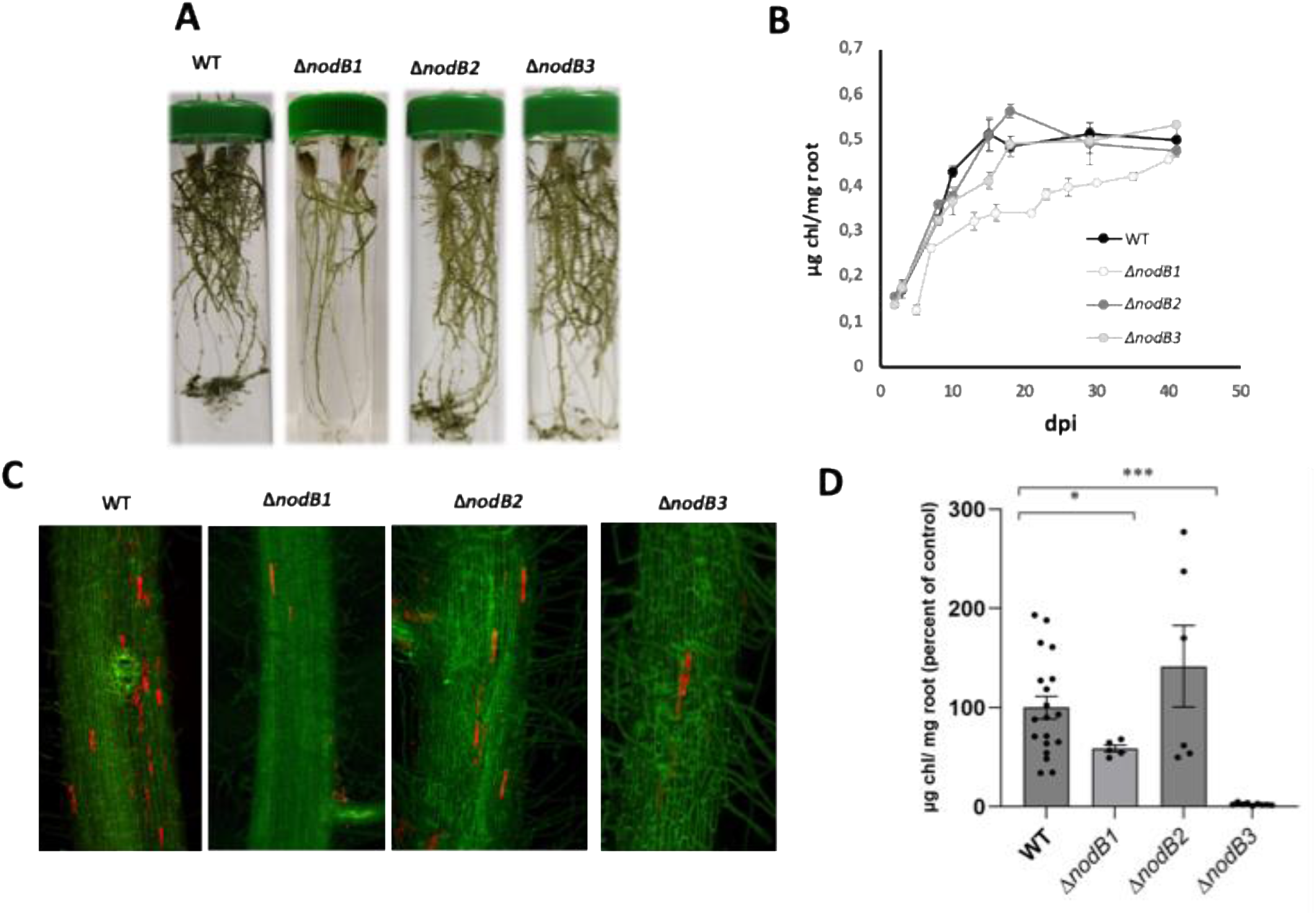
Biofilm formation and colonization of *O. sativa* roots by *N. punctiforme* and *nodB* deletion mutants. (A) Representative images showing biofilm formation on *O. sativa* roots at 15 days post inoculation (dpi). (B) Kinetics of biofilm formation quantified as chlorophyll *a* content per mg of root over time. Values are presented as mean ± standard error of the mean (SEM). (C) *O. sativa* roots inoculated with *N. punctiforme* or *nodB* deletion mutants were visualized at 45 dpi by confocal laser scanning microscopy. Z-stacks were generated from 90 to 130 optical sections. Plant cell wall autofluorescence is shown in green, while cyanobacterial chlorophyll autofluorescence is shown in red. Scale bar = 150 µm. (D) Quantification of endophytic colonization expressed as chlorophyll *a* content per mg of root. Data were normalized to their respective controls, defined as 100%. Values are presented as mean ± SEM. Statistical significance is indicated as: * p < 0.05; *** p < 0.001.

Symbiotic competence was further assessed by quantifying plant colonization (Fig. 3). The Δ*nodB1* mutant showed colonization levels approximately 40% lower than the wild type, indicating that its early defect in attachment partially affects the colonization phenotype. In contrast, the Δ*nodB2* mutant showed no reduction in colonization efficiency compared to the wild type, although the high variability of the data prevented robust statistical conclusions (P > 0.05). Strikingly, despite no significant differences being observed in root association compared to the wild type, the Δ*nodB3* mutant exhibited a severe impairment in colonization, with values close to background levels, representing a reduction of approximately 98% compared to the wild type (P < 0.001).

### 3.4. NodD regulators differentially control symbiotic competence in *N. punctiforme*

Distinct phenotypic differences were observed among the *nodD* mutants. The *nodD1::pRL278* mutant and Δ*nodD2* mutants did not show significant deviations from the wild-type strain in filament morphology, cellular structure, or growth. In contrast, the Δ*nodD3* mutant exhibited clear alterations in cellular and filament morphology, including enlarged cells and filaments with impaired differentiation into hormogonia, although heterocyst formation was still observed. In addition, the Δ*nodD3* mutant showed a significantly reduced growth rate compared to the wild type, both in solid and liquid media (Fig. S2; Fig. 4; Table 3).

**Figure 4.**
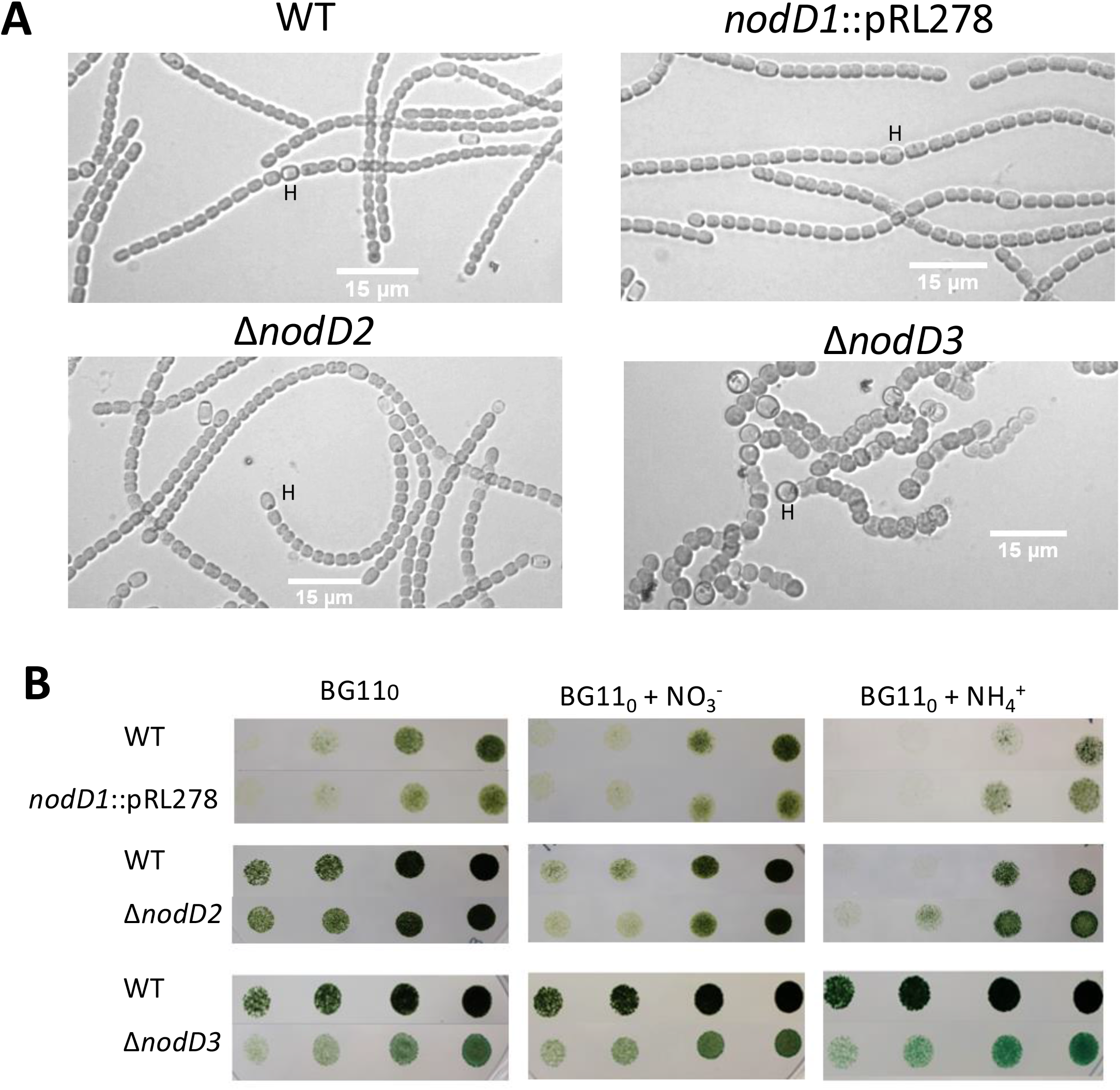
Morphological and growth characterization of *Nostoc punctiforme* WT and *nodD* deletion mutants. (A) Bright-field microscopy images (40× objective) of WT and *nodD1* :: pRL278, Δ*nodD2*, and Δ*nodD3* strains grown in BG110 medium. Vegetative cells and heterocysts are visible; representative heterocysts are indicated with “H”. (B) Spot assay of WT and mutant strains after 9 days of growth on solid media. Serial drops containing 5, 10, 25, and 50 ng of chlorophyll *a* (left to right) were spotted on BG110 and BG110 plates supplemented with combined nitrogen sources, as indicated.

**Table 3.** Morphological characterization of the wild-type (WT) strain and the. ΔnodD or mutants of *N. punctiforme* grown under diazotrophic conditions. Filament length (number of vegetative cells per filament), heterocyst frequency (% of total cells), vegetative cell length (µm), vegetative cell width (µm), and cell aspect ratio (length/width) are presented as mean ± standard error of the mean (SEM). Hormogonia formation was assessed qualitatively after induction and is indicated as present (+) or absent (−). Quantifications were performed from approximately 100 filaments and 130 cells per condition. Statistical significance is indicated as: ^a^ p < 0.05; ^b^ p < 0.001.

| Strain | Filament length (cells filament<sup>-1</sup>) | Heterocyst percentage (%) | Cell Length (μm) | Cell width (μm) | Aspect ratio (L/W) | Hormogonia formation |
| --- | --- | --- | --- | --- | --- | --- |
| WT | 22.21 ± 0.89 | 6.90 ± 0.38 | 2.50 ± 0.06 | 2.17 ± 0.02 | 1.17 ± 0.03 | + |
| *nodD1::pRL278* | 21.05 ± 0.80 | 7.88 ± 0.52 | 2.81 ± 0.06<sup>b</sup> | 2.45 ± 0.02<sup>b</sup> | 1.16 ± 0.03 | + |
| *ΔnodD2* | 25.53 ± 1.10<sup>a</sup> | 6.98 ± 0.38 | 2.49 ± 0.04 | 2.31 ± 0.02<sup>b</sup> | 1.09 ± 0.02<sup>a</sup> | + |
| *ΔnodD3* | 14.54 ± 0.80<sup>b</sup> | 12.1 ± 0.70<sup>b</sup> | 2.14 ± 0.06<sup>b</sup> | 2.99 ± 0.03<sup>b</sup> | 0.72 ± 0.02<sup>b</sup> | - |

To evaluate the role of NodD regulators during plant interaction, the kinetics of association to *O. sativa* roots were analysed over time. At early time points (3 days post-inoculation, dpi), the Δ*nodD3* mutant displayed a consistent and significant reduction in root association compared to the wild type (0.119 ± 0.013 vs 0.173 ± 0.006; ∼31% decrease; P < 0.01) (Fig. 4). Similarly, the Δ*nodD2* mutant showed a moderate but significant decrease in early association levels at 3 dpi (0.147 ± 0.009 vs 0.173 ± 0.006; ∼15% reduction; *P* < 0.05). These reductions persisted up to 8 dpi, with Δ*nodD2* and Δ*nodD3* exhibiting significantly lower association levels than the wild type (0.289 ± 0.011 and 0.247 ± 0.014, respectively, compared with 0.324 ± 0.011 for the wild type), corresponding to reductions of ∼11% (*P* < 0.05) and ∼24% (*P* < 0.01). In contrast, the *nodD1::pRL278* mutant displayed association kinetics comparable to those of the wild type at all early time points, with no statistically significant differences detected (*P* > 0.05).

At later stages (days 10–40), differences among mutants persisted. While *nodD1::pRL278* mutant maintained association levels similar to the wild type throughout the experiment (e.g., day 15: 0.475 ± 0.015 vs 0.513 ± 0.037; P > 0.05), both Δ*nodD2* and Δ*nodD3* exhibited a marked and persistent impairment in root association. At 15 dpi, association values for both mutants were approximately 0.36 ± 0.013, representing a ∼29–30% reduction relative to the wild type (0.513 ± 0.037; *P* < 0.01). This defect remained stable throughout the experimental period and was still evident at 40 dpi, when reductions of approximately 32% and 34% were observed for Δ*nodD2* and Δ*nodD3*, respectively (Fig. 4).

Symbiotic competence was further assessed by quantifying plant colonization, with values normalized to the wild type, which was set to 100%. The *nodD1::pRL278* mutant showed colonization levels comparable to those of the wild type. In contrast, the Δ*nodD2* mutant displayed a significant reduction in colonization, with mean values of 4.83 ± 1.48 % of the wild type colonization level, corresponding to a decrease of approximately 95% (P < 0.001). Strikingly, the Δ*nodD3* mutant exhibited an even stronger colonization defect, with values close to background (0.22 ± 0.12 % of the wild type level), representing a reduction of more than 99% compared to the wild type (P < 0.001) (Fig. 5).

**Figure 5.**
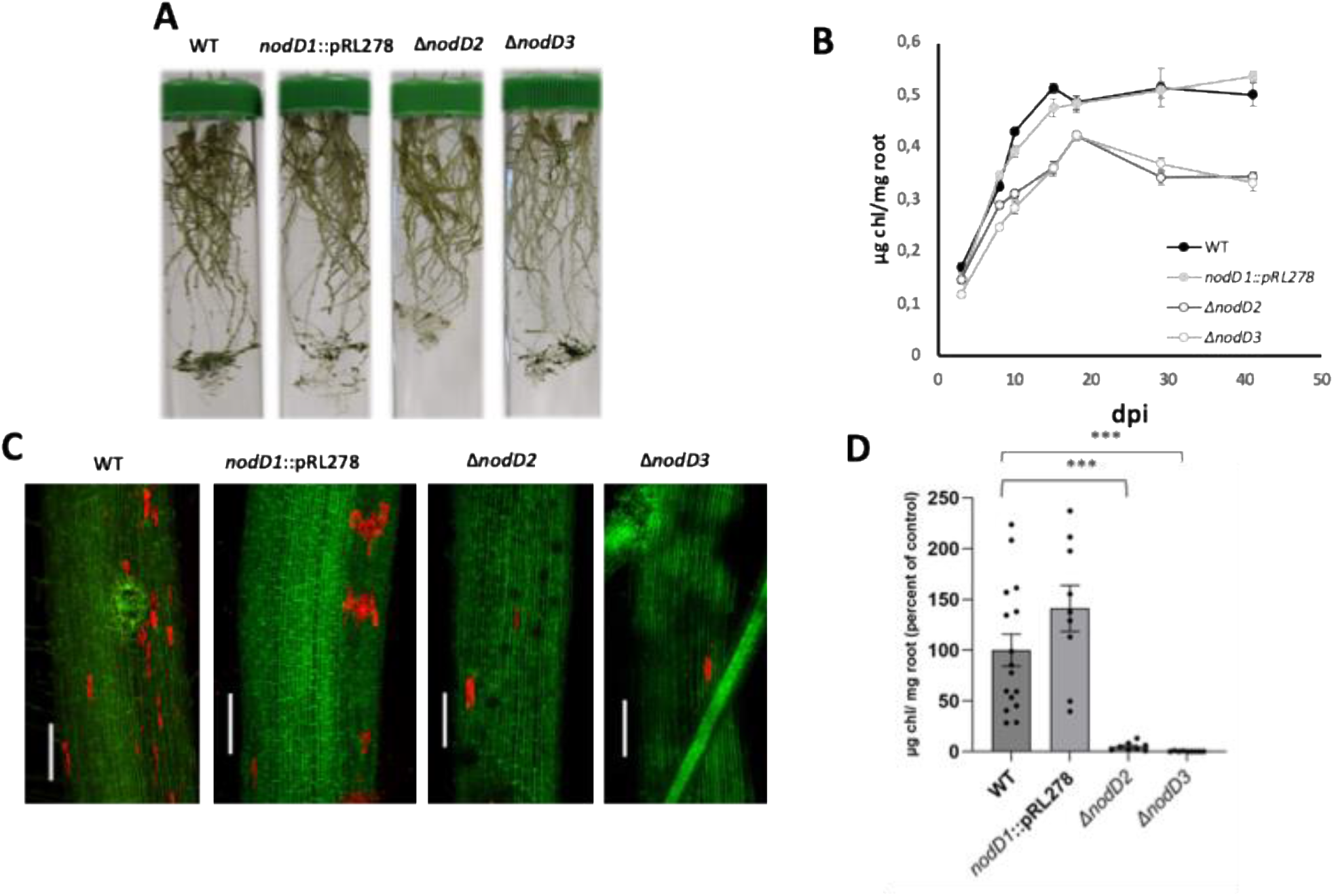
Biofilm formation and colonization of *O. sativa* roots by *N. punctiforme* and *nodD* mutants. (A) Representative images showing biofilm formation on *O. sativa* roots at 15 days post inoculation (dpi). (B) Kinetics of biofilm formation quantified as chlorophyll *a* content per mg of root over time. Values are presented as mean ± standard error of the mean (SEM). (C) *O. sativa* roots inoculated with *N. punctiforme* or *nodD* mutants were visualized at 45 dpi by confocal laser scanning microscopy. Z-stacks were generated from 90 to 130 optical sections. Plant cell wall autofluorescence is shown in green, while cyanobacterial chlorophyll autofluorescence is shown in red. Scale bar = 150 µm. (D) Quantification of endophytic colonization expressed as chlorophyll *a* content per mg of root. Data were normalized to their respective controls, defined as 100%. Values are presented as mean ± SEM. Statistical significance is indicated as: * p < 0.05; *** p < 0.001.

## 4. DISCUSSION

This study provides functional evidence that Nod-like components in *N. punctiforme* play a key role in the establishment of symbiotic interactions with *O. sativa*. Rather than acting as general determinants of cyanobacterial physiology, NodB and NodD homologues appear to be specifically involved in regulating distinct stages of plant interaction, supporting the existence of a complex signalling and regulatory system analogous, but not identical, to the Nod factor pathway described in rhizobia.

These findings contribute to filling a major gap in our understanding of the molecular basis of plant–cyanobacterium symbiosis, a system that, despite its ecological and agronomic relevance, remains far less characterized than other plant–microbe associations. In particular, the identification of functional roles for Nod-like components provides a mechanistic entry point to investigate how cyanobacteria coordinate the transition from free-living to symbiotic lifestyles and how this process is perceived by the plant host. More broadly, our results reinforce the idea that evolutionarily distant microorganisms may converge on shared signalling strategies to engage plant symbiotic pathways, while retaining lineage-specific regulatory features.

### 4.1. Functional diversification of NodB homologues suggests a modular biosynthetic system

The absence of detectable phenotypic alterations in free-living conditions for all *nodB* mutants indicates that these genes are not required for core aspects of cyanobacterial physiology, including growth, morphology, and cellular differentiation under the conditions tested. This contrasts with many enzymes involved in primary metabolism and suggests that NodB homologues are not essential for basal cellular functions in *N. punctiforme*, but instead may be specifically associated with processes that are relevant during interaction with the plant host.

In rhizobia, NodB catalyses the deacetylation of the chitooligosaccharide backbone during lipochitooligosaccharide (LCO) biosynthesis, representing a key step in the generation of Nod factors (John et al., 1993; Mergaert & Van Montagu, 1997; Oldroyd, 2013). In this study, three candidate NodB homologues were selected in *N. punctiforme* based on structural similarity to rhizobial NodB proteins, with the aim of assessing their potential contribution to symbiosis.

The phenotypic analysis of these mutants reveals clearly distinct outcomes. The Δ*nodB2* strain does not exhibit detectable differences in growth, root association, or colonization, indicating that this protein is not required for symbiosis under the conditions tested. In contrast, both Δ*nodB1* and Δ*nodB3* display symbiosis-related phenotypes. The Δ*nodB1* mutant shows reduced and delayed attachment together with a moderate reduction in colonization, suggesting a contribution to early stages of host interaction. More notably, the Δ*nodB3* mutant exhibits a severe defect in colonization despite normal association levels, pointing to a critical role at later stages of the symbiotic process. Interestingly, structural analysis revealed that NodB3 displays the highest TM-score among the NodB homologues, indicating a closer global structural similarity to rhizobial NodB proteins, which is consistent with its stronger functional impact on plant colonization.

This context-dependent phenotype suggests that the contribution of NodB homologues might be specifically required during host interaction, rather than under standard culture conditions. Such behaviour is consistent with regulatory or metabolic pathways that are induced by plant-derived signals or by the physiological changes associated with symbiotic transitions, such as hormogonia formation, surface recognition, or biofilm establishment. In this framework, genes that are dispensable for free-living growth may become critical determinants of symbiotic competence, highlighting the importance of analysing candidate functions under biologically relevant interaction conditions.

This interpretation is further supported by previous proteomic analyses of the early stages of the interaction, in which a NodB-like protein corresponding to NodB1 (Npun_F3876; B2J5A3) was found to be specifically induced in response to the plant partner (Álvarez et al., 2022). The induction of this protein during pre-symbiotic stages, together with the attachment and colonization defects observed in the Δ*nodB1* mutant, provides consistent evidence linking this enzyme to early steps of host interaction.

Taken together, these results indicate that not all NodB homologues contribute equally to symbiosis and identify NodB1 and, particularly, NodB3 as relevant components for host interaction. Given the conserved enzymatic role of NodB proteins in rhizobia, these candidates may be involved in the synthesis or modification of signalling molecules required for symbiosis, although the nature of these compounds remains unknown. Future work will be required to directly characterize the biochemical activity of these enzymes and to determine their specific substrates and functional roles in the context of cyanobacterial–plant interactions.

### 4.2. NodD homologues define a regulatory hierarchy controlling both symbiosis and cellular differentiation

In contrast to NodB, mutations in *nodD* homologues revealed a strong regulatory component, with clear differences in the magnitude and scope of the phenotypes observed. NodD1 did not show any phenotypic defect, that may be explained by the presence of WT copies in the genome, or because it is dispensable under the conditions tested. However, NodD2 and especially NodD3 might play critical roles in symbiotic competence, indicating the existence of a hierarchical regulatory organisation. These findings suggest that NodD homologues do not act redundantly, but rather contribute differentially to the coordination of the cyanobacterial response to the plant partner.

In rhizobia, NodD proteins function as LysR-type transcriptional regulators that activate nod gene expression in response to plant-derived flavonoids, thereby controlling LCO biosynthesis (Oldroyd, 2013; Mus et al., 2016). This regulatory framework establishes a direct mechanistic link between host signal perception and the production of symbiotic signals. Our results suggest that elements of this regulatory logic may be in *N. punctiforme*, but deployed within a more complex and expanded network. The moderate colonization defect observed in Δ*nodD2* in the absence of detectable alterations in growth or morphology, is consistent with a role in regulating symbiosis-specific functions without broadly affecting general physiology. This phenotype is compatible with a NodD-like function, operating downstream of plant signal perception, potentially controlling the expression of genes involved in early interaction processes, including signalling, adhesion, or the production of symbiotic factors. Analysis of these signalling network will be performed in future studies.

In contrast, the phenotype of Δ*nodD3* suggests a much broader regulatory role. This mutant is not only impaired in symbiotic competence but also displays defects in growth and cellular differentiation, including reduced or altered hormogonia formation. Given that hormogonia are the specialized infection units required for host colonization in cyanobacteria (Meeks & Elhai, 2002) these observations stablish a connection between NodD3 function and the developmental transitions underlying symbiotic competence. This suggests that, unlike canonical NodD regulators in rhizobia, NodD3 may integrate environmental sensing with developmental regulation, coordinating the physiological switch from a free-living to a symbiotic state. This model is further supported by structural analysis, as NodD3 displays the highest TM-score among NodD homologues, indicating a strong conservation of global protein architecture relative to rhizobial NodD regulators (Xu and Zhang, 2010). Such structural conservation is consistent with its broad phenotypic impact and supports its role as a key regulatory component linking environmental sensing, cellular differentiation, and symbiotic competence. Such integration is likely to be particularly relevant in cyanobacteria, where symbiosis requires extensive reprogramming of cellular behaviour, including motility, surface attachment, and metabolic adaptation (Meeks & Elhai, 2002; Álvarez et al., 2023). In this context, NodD3 may function as a higher-level regulator controlling multiple downstream pathways required for successful host interaction. Rather than acting solely as an activator of signal biosynthesis genes, it may coordinate a broader regulatory programme linking signal perception, cellular differentiation, and infection competence.

The coexistence of multiple NodD homologues with distinct phenotypic outputs therefore points to a regulatory network rather than a single linear pathway. This architecture may provide *N. punctiforme* with the flexibility required to respond to diverse environmental cues and host-derived signals. Such regulatory plasticity is consistent with the ecological versatility of this cyanobacterium, which is capable of establishing symbioses with phylogenetically distant plant hosts. In this framework, NodD2 and NodD3 may represent functionally differentiated nodes within a broader regulatory hierarchy, contributing in a coordinated manner to the establishment and progression of the symbiotic interaction.

### 4.3. A working model for Nod-like signalling in cyanobacterial symbiosis

The combined genetic and phenotypic evidence presented in this study supports a model in which Nod-like components in *N. punctiforme* define a coordinated signalling framework controlling distinct stages of plant interaction. Integration of the functional specialization observed among NodB homologues with the regulatory hierarchy mediated by NodD proteins suggests the existence of a multi-layered system linking signal production, environmental sensing, and developmental transitions.

In this model, NodD regulators act as key nodes integrating plant-derived or environmental cues, thereby controlling the expression of downstream genes required for symbiotic competence. NodD2 appears to regulate a subset of symbiosis-specific functions, primarily affecting early interaction and colonization efficiency, whereas NodD3 may function as a higher-order regulator coordinating broader physiological transitions, including hormogonia differentiation and growth. This regulatory architecture is consistent with the requirement for precise temporal control of infection-related processes in cyanobacteria. Future studies will aim to define the NodD2 and NodD3 regulons, with the goal of testing the proposed model and elucidating the regulatory networks underlying Nod-like signalling in *N. punctiforme*.

Concomitantly, NodB homologues may contribute to the biosynthesis or modification of signalling molecules required at different stages of the interaction. The distinct phenotypes of Δ*nodB1* and Δ*nodB3* suggest that these enzymes are not functionally redundant but instead participate in temporally or mechanistically differentiated steps of the symbiotic process, possibly influencing early recognition events and later colonization stages, respectively.

Together, these elements support a model in which *N. punctiforme* produces signalling molecules, potentially analogous to lipochitooligosaccharides, under the control of NodD-dependent regulatory pathways. These signals may be perceived by the host plant to activate the common symbiosis signalling pathway, thereby enabling intracellular accommodation. While the biochemical identity of these molecules remains to be determined, this framework provides a mechanistic basis for understanding how cyanobacteria coordinate the transition from free-living to symbiotic lifestyles. Future work should aim to identify the chemical nature of the signalling compounds involved and to define the regulatory networks controlled by NodD proteins. Such studies will be essential to validate this model and to determine how broadly conserved these mechanisms are across cyanobacterial lineages.

## CREDIT AUTHORSHIP CONTRIBUTION STATEMENT

C.A. and V.M. designed the research and the experimental setup, and were responsible for writing the manuscript. J.E.F. carried out the construction of the plasmids. C.S.S. and A.J.F generated the mutant strains. C.S.S. conducted the genetic analysis of the cyanobacterial strains and overall characterization. L.J.R. participated in the phenotypic characterization of the mutants. All co-authors reviewed, edited for clarity, and approved the final manuscript.

## DATA AVAILABILITY

The data generated in this work will be made available on request.

---

**Figure S1. PCR validation of the generated *Nostoc punctiforme* mutants.** Genomic DNA was used as template to verify the replacement of target genes by the Ω*npt* resistance cassette in deletion mutants (Δ*nodB1*, Δ*nodB2*, Δ*nodB3*, Δ*nodD2*, and Δ*nodD3*), as well as the integration of the suicide plasmid pRL278 in the *nodD1*::pRL278 insertion mutant. For deletion mutants, two types of PCR reactions were performed. Reactions 1 and 2 used primer combinations consisting of a gene-flanking primer (upstream or downstream of the target locus) paired with a primer annealing within the coding sequence of the target gene, and were used to detect the presence of the wild-type allele. Reactions 3 and 4 used primer combinations consisting of the same gene-flanking primers paired with primers annealing within the Ω*npt* cassette, and were used to confirm cassette insertion at the target locus. For the *nodD1*::pRL278 insertion mutant, primers flanking the coding region were used to detect the wild-type allele (reaction 5), whereas primer combinations including a flanking primer and a primer annealing within pRL278 were used to verify plasmid integration (reactions 6 and 7). Wild-type genomic DNA was used as a positive control for reactions 1, 2, and 5, and as a negative control for reactions 3, 4, 6, and 7. Expected amplicon sizes (bp) are indicated in each schematic representation. M, mutant genomic DNA; W, wild-type genomic DNA.

**Figure S2. Growth characterization of *Nostoc punctiforme* WT and** Δ**nodD3 mutant.** (A) Growth curves of WT and Δ*nodD3* strains grown in BG110 liquid medium under standard growth conditions using a multicultivator system as described in Materials and Methods. Data are shown as mean ± SEM. (B) Growth rates calculated from the exponential phase of the growth curves shown in (A), obtained by linear regression of log-transformed growth data. Data are shown as mean ± SEM.

**Table S1.**
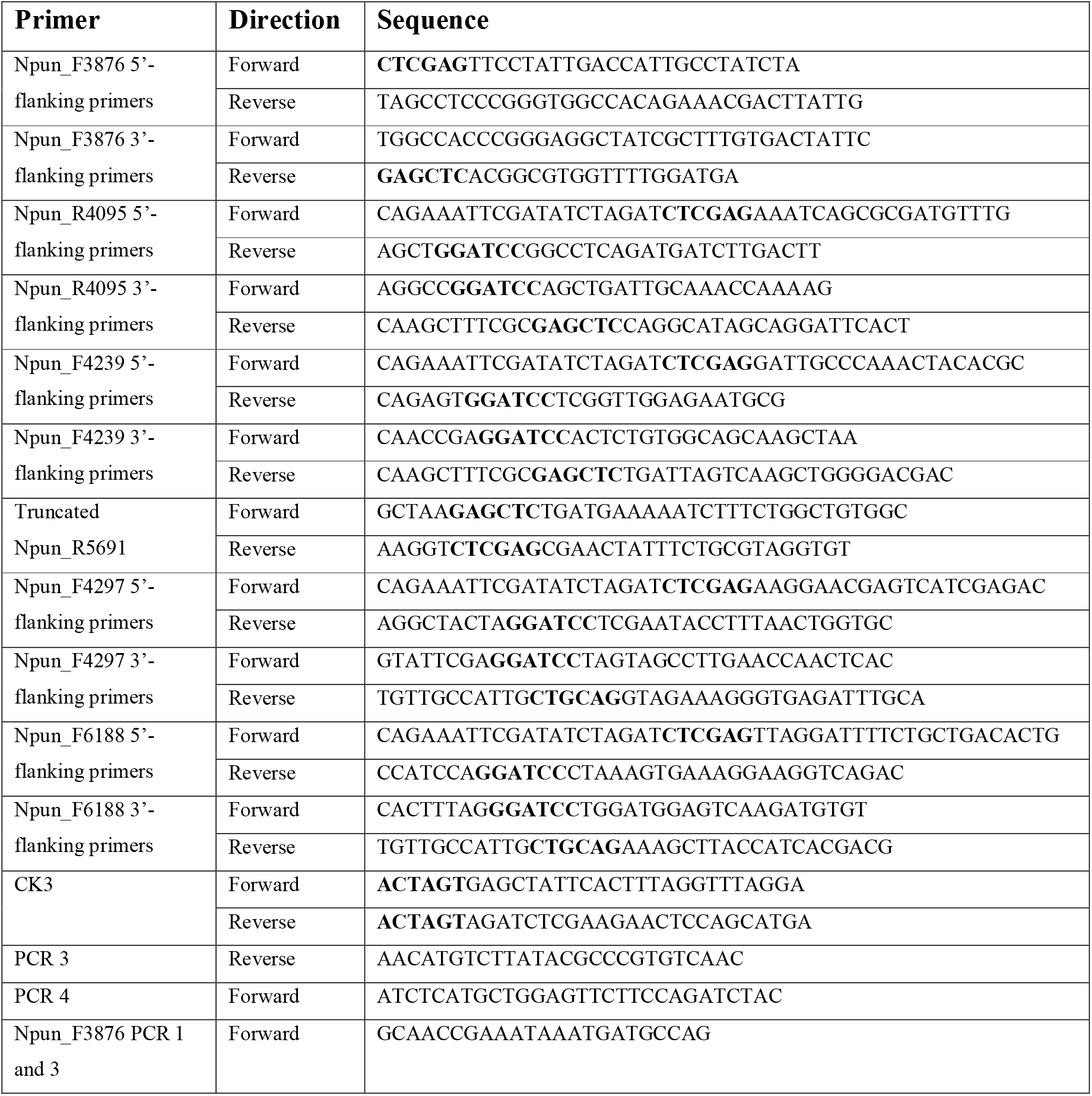
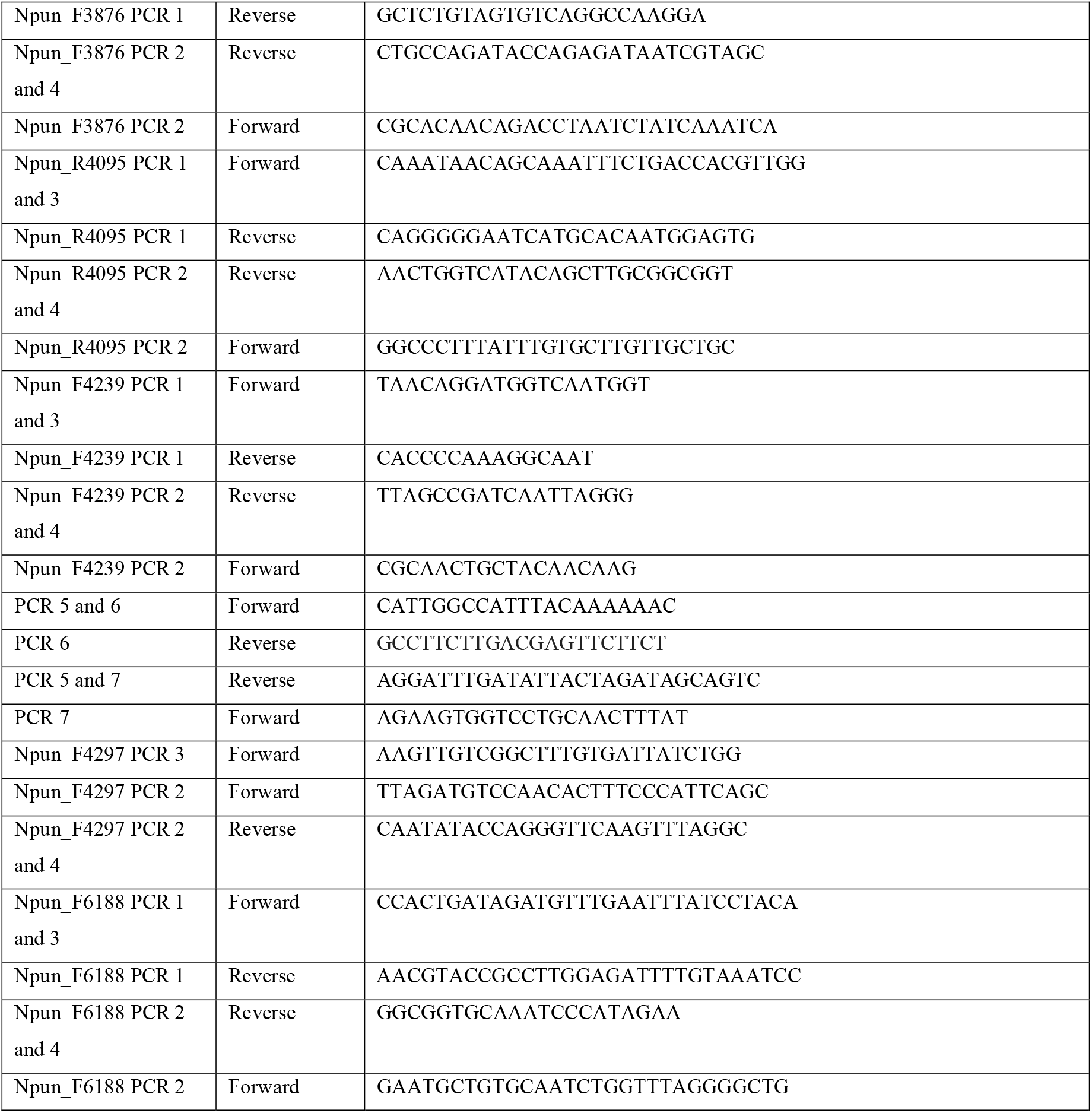
Oligonucleotide primers used in this study. Restriction endonuclease sites incorporated are shown in bold.

